# Serodiagnostic potential of synthetic peptides derived from African swine fever virus putative protein pCP312R

**DOI:** 10.1101/512772

**Authors:** Sylvester Ochwo, David Kalenzi Atuhaire, Mathias Afayoa, Majid Kisseka, Phillip Kimuda Magambo, Christian Ndekezi, Julius Boniface Okuni, William Olaho-Mukani, Lonzy Ojok

**Affiliations:** College of Veterinary Medicine, Animal resources and Biosecurity, Makerere University, P.O.BOX 7062, Kampala, Uganda; African Union-Inter African Bureau for Animal Resources, P.O.BOX 30786, Nairobi, Kenya

**Keywords:** African swine fever virus, Synthetic Peptides, Immunohistochemistry, ELISA, Serodiagnosis, Insilco prediction

## Abstract

African swine fever (ASF) is a hemorrhagic disease of domestic swine, with often high mortality rates registered. To date there is still no vaccine produced against ASF, and disease management in countries including Uganda, where the disease is endemic is dependent on accurate and timely diagnosis programs and quarantine. This study aimed at contributing more knowledge towards ASF diagnosis by investigating the serodiagnostic potential of synthetic peptides of an ASF putative protein pCP312R. Antigenic regions of the pCP312R putative protein were identified using Kolaskar and Tongaonkar antigenicity prediction method and twelve (12) peptides were predicted, out of which four (4) peptides were selected and synthesised. An additional peptide derived from the carboxyl end of the ASFV p54 protein was also synthesised and used as a control. Polyclonal rabbit antibodies raised against each of the five peptides was used in immunohistochemistry, and each demonstrated ability to localize viral antigen in pig tissue albeit with slightly varying intensities, at a dilution of 1:200, with antibodies against peptides cpr1, cpr2, cpr3 and cpr4 all accurately staining infected macrophages. However all the peptides evaluated in this study performed moderately when used in indirect ELISA tests giving the following results; CP1; diagnostic sensitivity of 55% (95% CI, 0.3421-0.7418) and specificity of 96% (95% CI, 0.8046-0.9929), CP2; diagnostic sensitivity of 100% (95% CI, 0.8389-1) and specificity of 52% (95% CI, 0.335-0.6997), CP3; diagnostic sensitivity of 95% (95% CI, 76.39-99.11) and specificity of 88% (95% CI, 70.04-95.83), CP4; diagnostic sensitivity of 90% (95% CI: 0.699-0.9721) and specificity of 76% (95% CI: 0.5657-0.885) and p54; diagnostic sensitivity of 100% (95% CI, 0.8389-1) and specificity of 56% (95% CI, 0.3707-0.7333). This study presents the first time synthetic peptides have been successfully predicted, designed and evaluated for Serodiagnosis of African swine fever in domestic pigs. This study in addition showed that there is potential for use of polyclonal anti-peptide antibodies in the diagnosis of ASF using immunohistochemistry.

## Background

African swine fever (ASF) is an extremely lethal hemorrhagic disease of domestic swine, with mortality rates approaching 100%. ASF occurs in several disease forms, ranging from highly lethal to sub clinical infections depending on contributing viral and host factors (Zsak *et al*., 2005).

The causative agent, African Swine Fever Virus (ASFV), is a unique and genetically complex DNA virus. This virus is large and icosahedral in shape, with linear double stranded DNA genome ranging in size between 170 kilo base pairs (kbp) and 190 kbp, and encoding approximately 165 viral proteins (Andrés *et al*., 2002). ASFV is the only member of the family Asfarviridae and the only known DNA arthropod borne virus (arbovirus) (Dixon *et al*., 2004). In eastern and southern Africa, cycling of ASFV between soft ticks of the genus Ornithodoros and warthogs (Phacochoerus africanus), provides a natural reservoir for the virus, that poses a continuous threat to domestic pig populations globally (Anderson *et al*., 1998). In addition to this, the presence of asymptomatic animals due to occurrence of virus strains with low virulence makes serological diagnosis essential for the control of the disease (ASF) in countries affected. The continuing occurrence and further spread of ASF has hampered pig sector development in sub-Saharan Africa, leaving it well below its full potential (FAO, 2012).

At present there is still no vaccine against ASF. Management of the disease in countries where the disease is endemic is only based on proficient diagnosis programs, quarantine and by sacrificing infected animals. Diagnostic techniques for ASF are focused on antigen, antibody, and genomic DNA detection by Polymerase Chain Reaction (PCR), and on virus isolation and localization in clinical samples. Field diagnosis of ASF is confirmed in the laboratory by virus isolation, since the clinical signs are not confirmatory for ASF. These available diagnostic techniques have varying levels of cost per test, suitability and also vary in sensitivity and specificity (Oura *et al*., 2012). When the virus strains that cause the disease are highly virulent and kill pigs before they mount an antibody response, PCR is used as an important tool for ASFV diagnosis. However, because of the existence of strains of reduced virulence that result in a low mortality (Bech-Nielsen *et al*., 1995), ASF is diagnosed mainly by detection of specific antibodies. Therefore, it is important to study the viral components that are capable of inducing humoral immune responses and are therefore suitable for use as diagnostic reagents (Gallardo *et al*., 2006).

In this study we investigated the immunogenicity and diagnostic potential of synthetic peptides derived from antigenic regions of the ASFV putative protein pCP312R, encoded by the Open reading frame (ORF) CP312R. This protein was earlier identified by screening an ASFV expression library with hyper immune porcine serum from an infected domestic pig (Kollnberger *et al*., 2002). This suggested that this protein may be useful in the serodiagnosis of ASF. The serodiagnostic potential of a peptide sequence from ASFV protein p54 which is known to be highly immunogenic (Barderas *et al*., 2001; Rodriguez *et al*., 1994) was also analysed in this study and used as a control.

## Methodology

### Study sites

The study followed a Laboratory experimental design, and was performed at the Molecular Biology Laboratory (MOBILA), Department of Bio-molecular Resources and Bio-lab sciences at the College of Veterinary Medicine, Animal Resources and Bio-security, Makerere University, Kampala.

### Materials used in the study

Synthetic peptides were selected using *in silico* techniques (Kolaskar & Tongaonkar, 1990) and synthesised by Thermo-Scientific, Germany. For each selected peptide, a conjugated and a none conjugated version were synthesised. New Zealand white rabbits, 7 months old were used in immunisation/immunogenicity studies. Tissue samples collected from dead pigs that had been experimentally infected with ASFV were used for DNA extraction and for ASF diagnostic PCR to confirm ASF infection. The tissues were subsequently fixed in 10% buffered formalin and later used for immunohistochemistry.

### Extraction of viral DNA from tissue samples

Tissues (spleen, lung, liver and lymph nodes) were collected from dead pigs that had been experimentally infected with ASFV genotype IX (Afayoa *et al*., 2014) and DNA was extracted using the DNeasy blood and tissue kit Qiagen. This was then stored at −80°C until used.

### Amplification of viral DNA by PCR

ASF diagnostic primers described by (Aguero *et al*., 2003) were used to amplify a 257 bp region corresponding to the central portion of the p72 gene, thus confirming presence of ASFV DNA in a tissue sample. The primer sequences used were as follows: Primer PPA-1 sequence 5’-AGT-TAT-GGG-AAA-CCC-GAC-CC-3’ (forward primer); primer PPA-2 sequence 5’-CCC-TGA-ATC-GGA-GCA-TCC-T-3’ (reverse primer).

### Peptide design and selection

The ASFV putative gene and protein sequences were obtained from the NCBI database. This was then used for peptide selection using a semi-empirical method, which predicts antigenic determinants on proteins by using physicochemical properties of amino acid residues together with their frequencies of occurrence in experimentally known segmental epitopes, as described by (Kolaskar & Tongaonkar, 1990). Before selection of antigenic epitopes the pCP312R protein sequence was compared to other similar sequences available in the GenBank (Fig: 1).

**Fig 1:**
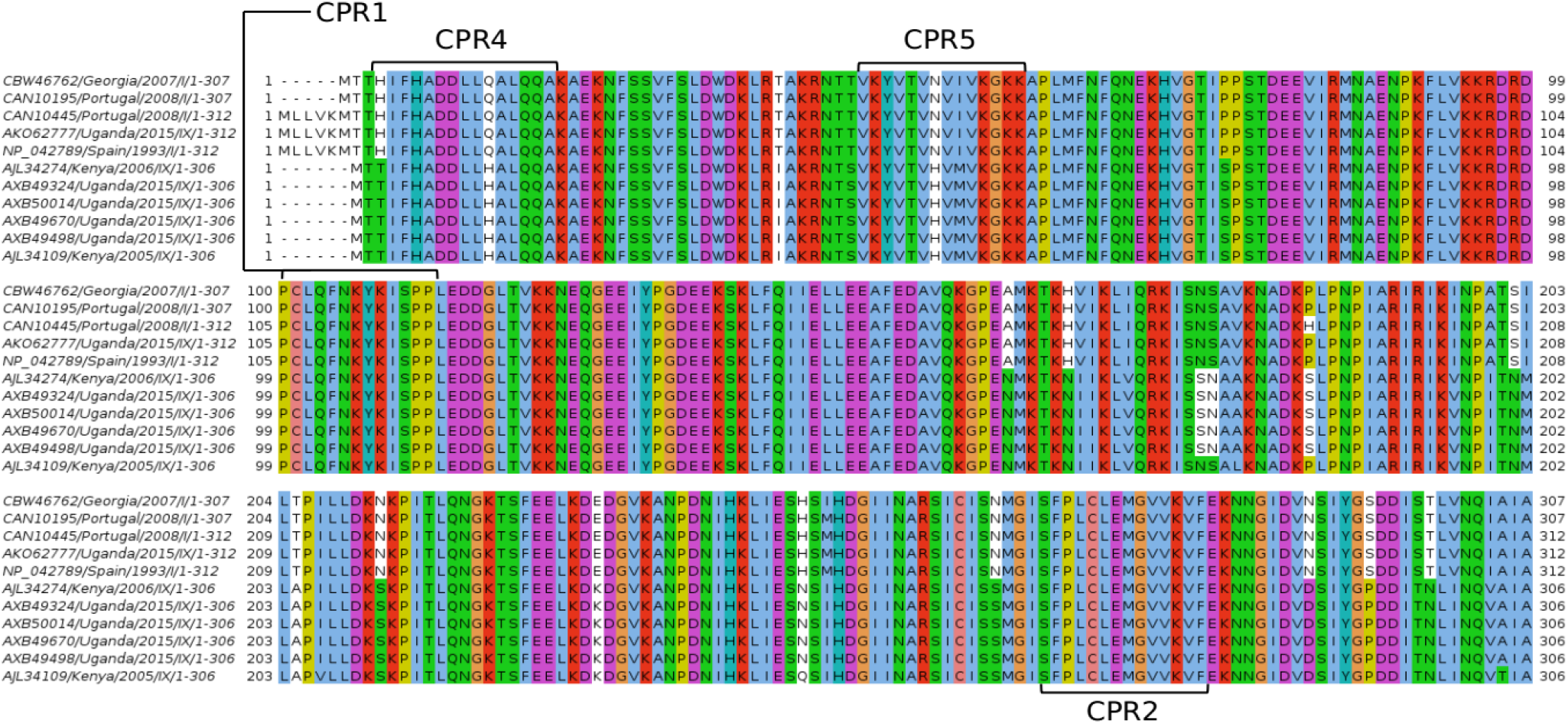
A multiple sequence alignment showing the amino acid sequence similarity between the pCP312R protein sequences used in his study. There is a high level of homology in the pCP312R protein sequences from different ASFV genotypes. The regions used for peptide selection have been marked CPR1, CPR2, CPR4 and CPR5.

The pCP312R protein sequence was then copied and pasted on to the analysis window of the prediction software and then the query submitted. Following this a list of antigenic determinants (peptides) was generated showing the peptide number, start and stop position on the protein sequence and length of each peptide. Four peptide sequences were then selected based on their length and position, which is those with 14 residues or more and found on the amino end, middle and carboxyl end of the pCP312R sequence.

The selected peptide sequences were then sent to Thermo scientific Germany for synthesis using Fmoc solid-phase technology and then purified using HPLC. During synthesis, two versions of each peptide were synthesised, a conjugated and a none conjugated version.

The peptides were conjugated at C-termini or cysteine side chains to Imject Mariculture Keyhole Limpet Haemocyanin (mcKLH) using carboxyl-reactive EDC cross linking or sulfhydryl-reactive maleimide chemistries respectively. After the coupling reaction the protein-peptide conjugates were then dialyzed to remove non-conjugated peptides and salts thus providing the highest possible immunogenicity.

### ASF p54 peptide selection

The ASF p54 synthetic peptide **CENLRQRNTYTHKDLENS** (residues 166 to 182), both un-conjugated and conjugated to mcKLH was selected for use because it has previously been described by (Rodríguez *et al*., 2004), where polyclonal antibodies were raised against the synthetic peptide, in rats. In this study this synthetic peptide was used to act a control during immunogenicity studies in rabbits and in addition its diagnostic potential was also investigated.

### Reconstitution of peptides

The lyophilised peptides were allowed to come to room temperature. Using information on the synthesis report provided by the manufacturer, reconstitution was done by adding the appropriate amount of 1X PBS pH 7.4 to each tube. This was followed by gently mixing to allow the lyophilisate to dissolve. Aliquots of 200μl each were then made off from each of the stock tubes and stored at −20°C for subsequent use in immunisation and ELISA.

### Immunisation of rabbits with selected peptides

Eight months old, healthy female New Zealand white rabbits not previously used in any experiment, were used in this immunisation study. They were divided into five (5) groups of two (2) rabbits each, named group one to five. Prior to immunisation each rabbit was bled and pre immune serum collected.

In preparation for immunisation, reconstituted aliquots of peptides conjugated to mcKLH were brought to room temperature then the required dose calculated. The appropriate volume was pipetted and transferred to a separate 1.5ml sterile tube. To this new tube 1X PBS was added to make a final volume of 400μl, followed by mixing. Two sets of adjuvant were then added, 100μl of 10mg/ml Quil A followed by 500μl of Alhydrogel to give a final volume of 1 ml. This was followed by mixing at ambient temperate, for 30 minutes on a rotary shaker.

The amount of conjugated peptide used for immunisation was 200μg of peptide per animal for the primary immunisation, and 100μg per animal for booster doses. Boosting was done on the 14th, 28th, and 42nd day after the primary immunisation. A total of three booster doses were administered. The subcutaneous route was used for immunisation, with the dose administered at four sites around the animals’ shoulder blades. Each animal group was immunised with a different peptide, as shown in the table 1;

**Table 1:**
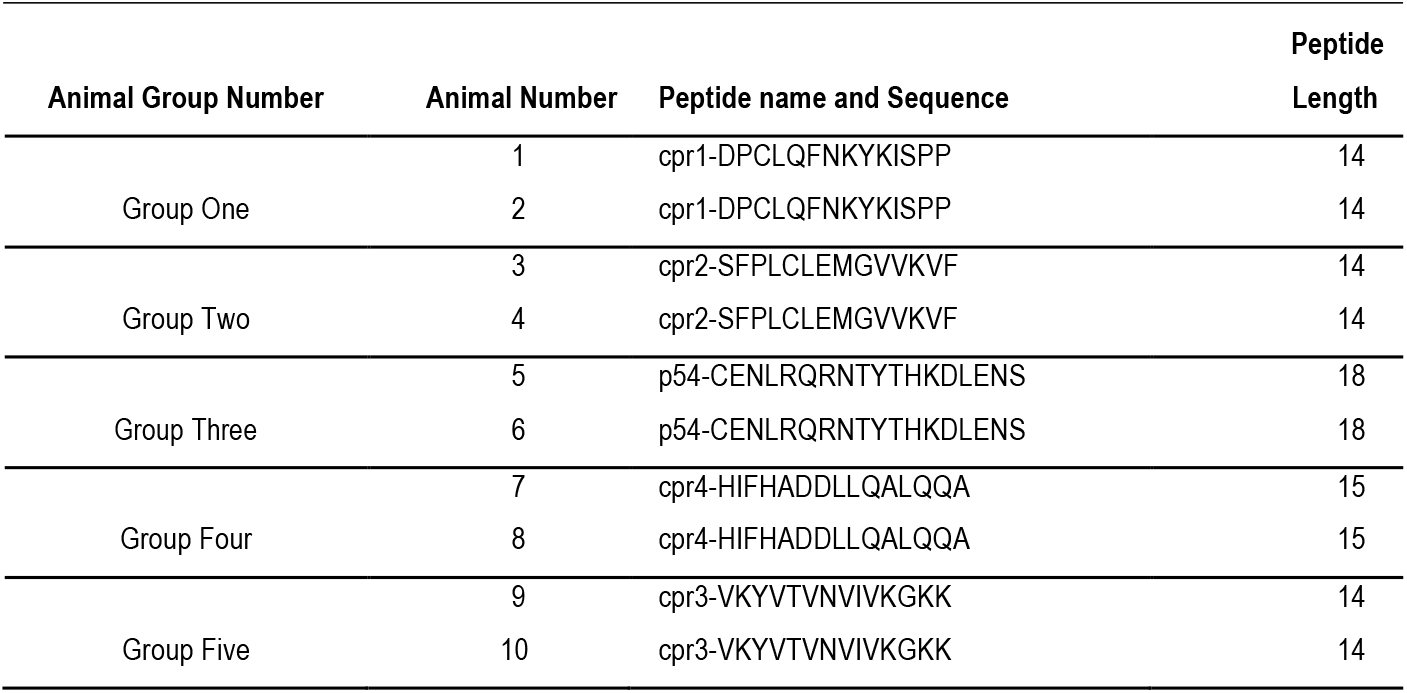
Shows a summary of the animal groups and corresponding peptide sequences used for the immunisation study

| Animal Group Number | Animal Number | Peptide name and Sequence | Peptide Length |
| --- | --- | --- | --- |
| Group One | 1 | cpr1-DPCLQFNKYKISPP | 14 |
|  | 2 | cpr1-DPCLQFNKYKISPP | 14 |
| Group Two | 3 | cpr2-SFPLCLEMGVVKVF | 14 |
|  | 4 | cpr2-SFPLCLEMGVVKVF | 14 |
| Group Three | 5 | p54-CENLRQRNTYTHKDLENS | 18 |
|  | 6 | p54-CENLRQRNTYTHKDLENS | 18 |
| Group Four | 7 | cpr4-HIFHADDLLQALQQA | 15 |
|  | 8 | cpr4-HIFHADDLLQALQQA | 15 |
| Group Five | 9 | cpr3-VKYVTNVIVKGKK | 14 |
|  | 10 | cpr3-VKYVTNVIVKGKK | 14 |

### Collection and processing of sera

Pre-immune serum was collected from each rabbit belonging to the five groups, by bleeding through the marginal ear vein; this was done before immunisation. The area on the ear was shaved with a number 21 size scalpel blade and disinfected with 70% ethanol before making a small slit on the marginal ear vein and allowing blood to drop slowly into an 8.5ml collection tube free from anticoagulant. This serum was used as the first negative control for comparison with immune sera. For collection of immune serum, bleeding of all five groups of rabbits was done on every 5th and 10th day after immunisation/booster dose using the same procedure described above.

After collection of blood in tubes, the blood was left to clot at room temperature for 45 minutes and then transferred to a refrigerator at 4°c overnight to allow maximum retraction of the clot. The blood was then centrifuged at 3000rpm for 10 minutes to separate serum from blood cells. This serum was then drawn into 1.8ml serum tubes, labeled and stored at −20°c until when required for analysis.

### Determination of antibody titers from collected rabbit sera

This was determined by use of an indirect ELISA. All the sera collected during the different stages of the immunisation study i.e. pre immune and immune sera after the different boosts, were subjected to analysis by ELISA to monitor immunogenicity of individual synthetic peptides.

The wells of an ELISA plate (Nunc, Germany) were coated with 100ng per well of un conjugated peptide in 100μl of 0.05M bi-carbonate coating buffer (pH 9.6) overnight at 4°C. Each peptide was used to coat two columns of the ELISA plate and the last two columns of the plate only received coating buffer with no peptide. The following day excess coating buffer was poured out of the plate, then the plate was blocked with 200μl per well of blocking buffer containing 5% Skimmed milk in 1X PBS, for 2 hours at room temperature.

Primary antibody (sera) collected from the five groups of rabbits was then prepared by diluting 1:20,000 in 1X PBS containing 1% skimmed milk. Serum from each rabbit group was then added to wells coated with the corresponding peptide, the volume added was 100μl of the diluted (1:20,000) serum. The wells in the last two columns of the plate received buffer with no serum. The plate was then incubated for 2 hours at room temperature. The plate was there after washed 6 times in washing buffer (PBS-T + 0.05% Tween20). 100μl per well of secondary antibody (Goat anti-rabbit IgG conjugated with Horse-raddish peroxidase) at a dilution of 1:5000 in dilution buffer was then added and the plate incubated at 37° C for 1 hour. The plate was washed 3 times with washing buffer and 100μl per well of one step 3,3’,5,5’-Tetramethylbenzidine (TMB) substrate (Thermo scientific, Germany) added and incubated for 15 minutes at room temperature. The reaction was then stopped with 100μl per well of stop solution (1M Sulphuric acid) and optical densities (OD) measured at 450 nm with a microplate reader (BioTek, United States).

### Evaluation of diagnostic potential of polyclonal anti peptide rabbit sera by immunohistochemistry

Polyclonal sera collected from the five groups of rabbits was subjected to immunohistochemical analysis using tissue from pigs confirmed to be ASF positive by OIE PCR. Paraffin embedded sections were dewaxed in three changes of xylene for 5 minutes each and then through three changes of absolute alcohol for 2 minutes each, before finally bringing the sections to distilled water.

The sections were then treated in sodium citrate buffer pH 6.0 by boiling for 10 minutes and then left to cool in the same buffer for 20 minutes, then washed with two changes of PBS. Sections were then washed in two changes of 1X PBS for 5 minutes each. Endogenous peroxidase was blocked using 3% hydrogen peroxide in methanol, followed by two 5 minute washes in 1X PBS. Non-specific binding was blocked with normal Goat serum (1:10 in 1X PBS) for 30 minutes.

Checkerboard titrations were done to obtain optimal dilutions of antibodies. The primary antibody (100μl of rabbit serum at a dilution of 1:200) was added to each section and incubated for an hour at 37 °C; two negative control slides received PBS and pre-immune serum while two positive control slides received anti-p54 peptide serum and mouse monoclonal anti-vp73 antibodies at a dilution of 1:400 (Cobbold *et al*., 1996). After incubation the slides were washed in two changes of PBS for 5 minutes each.

Secondary antibody, 100μl of goat anti-rabbit /anti mouse antibody conjugated to biotin (Histofine SAB-PO kit, Nichirei, Tokyo, Japan) was added to each of the slides and incubated for 10 minutes, followed by two 5 minute washes in PBS, 100ul of Horse raddish peroxidase enzyme reagent (Histofine SAB-PO kit, Nichirei, Tokyo, Japan) was then added to each section and incubated for 5 minutes. Two five minute washes in PBS were then done. This was followed by 100μl of 3, 3-diaminobenzidine tetrahydrochloride (DAB) substrate solution (Histofine SAB-PO kit, Nichirei, Tokyo, Japan) for five minutes. Following colour development the sections were washed with distilled water, then counterstained with haematoxylin (Sigma, Germany), for 40 seconds and differentiated in tap water for about 15 minutes. The sections were then dehydrated in three changes of absolute ethanol for two minutes each and cleared in three changes of Xylene for two minutes each before being permanently mounted with DPX.

### Evaluation of diagnostic potential pCP312R peptides by ELISA

Checkerboard titrations (CBT) were done for each of the peptides to determine the optimal concentrations and conditions for the ELISA, as described by (Crowther, 2000). This involved coating ELISA plates with varying concentrations of each peptide and titrating against varying concentrations of field sera (primary antibody) and secondary antibody (Goat anti-porcine IgG conjugated to HRP). Positive and negative field sera screened by an OIE recommended ELISA were selected for use in this CBT and were used to determine cut offs. The optimal conditions for the peptide ELISA were then determined and used to screen 45 porcine field serum samples against each of the 5 peptides. The serum samples used here were prescreened by OIE recommended ELISA and PCR and 20 were positive while 25 were negative by both ELISA and PCR respectively.

After optimisation, 5 plates were coated, each with a different peptide named CP1, CP2, CP3, CP4 and p54. This was done with 100μl per well of 4g/ml of un conjugated peptide in 0.05M bi-carbonate coating buffer (pH 9.6) overnight at 4°C. The following day excess coating buffer was poured out of the plate, then the plates were blocked with 200μl per well of blocking buffer containing 5% Skimmed milk in 1X PBS, for 2 hours at room temperature.

Primary antibodies (porcine field sera) were prepared for each of the plates by diluting the 45 field samples 1:4000 in 1X PBS containing 1% skimmed milk. Adopted positive and negative controls were added in duplicate to each plate, followed by the samples, the volume added was 100μl per well and loading was done in duplicate. The plates were then incubated for 2 hours at room temperature with agitation.

The plates were there after washed 6 times in washing buffer (0.05% PBS-Tween20). 100μl per well of secondary antibody (Goat anti-porcine IgG conjugated with Horse-raddish peroxidase) at a dilution of 1:10000 in dilution buffer was then added and the plate incubated at 37° C for 1 hour with no agitation. The plates were washed 3 times with washing buffer and 100μl per well of one step TMB substrate (Thermo scientific) added and incubated for 20 minutes at room temperature. The reaction was then stopped with 100μl per well of stop solution (1M Sulphuric acid) and ODs measured at 450 nm with a microplate reader (BioTek, United States).

Cut offs for positive samples were determined by using the formular: average OD of negative control x 3 standard deviations + OD of negative control. There after samples that were positive and negative by each peptide ELISA were selected and used to compute diagnostic sensitivity and specificity by comparing these results with those of the OIE reference ELISA test.

The computations were performed following the OIE terrestrial manual 2013 and the number of samples to be analysed was determined following table 1 shown in chapter1.1.5. Sensitivity and specificity of the test was calculated as shown in the table 2;

**Table 2:** Calculation of diagnostic sensitivity and specificity

|  |  | Reference test Results |  |
| --- | --- | --- | --- |
|  |  | Known positives | Known Negatives |
| Test Results | Positive | TP | FP |
|  | Negative | FN | TN |

Diagnostic sensitivity
$TP / (TP + FN)$
Diagnostic specificity
$TN / (TN + FP)$
**TP**=True Positive, **TN**=True Negative, **FP**=False Positive, **FN**=False Negative

### Data Analysis

The data obtained from the ELISA was analysed by Microsoft excel version 2007 and DaG_stat (Mackinnon, A. 2000) and presented in percentages and graphical format.

### Ethics statement

Full ethical clearance was obtained from the Uganda National Council for Science and Technology (UNCST) and the College of Veterinary Medicine, Animal Resources and Bio-security, Makerere University under reference number VAB/REC/11/110. Animal welfare and care was ensured in accordance with the international Guideline on Animal Welfare and Euthanasia. Any experimental animal in pain or moribund was immediately euthanized to relieve it from further suffering. Clean water and commercial animal feeds

## Results

### Prediction of highly antigenic regions on pCP312R sequence

Using the Kolaskar and Tongaonkar antigenicity prediction method, the ASFV putative protein pCP312R sequence (accession Q65180) was analysed Antigenicity analysis of the pCP312R sequence gave an average antigenic propensity of 1.017, while the minimum and maximum antigenic propensities obtained were 0.878 and 1.196 respectively. A total of 12 peptides were then predicted, with lengths carrying from 7 to 15 amino acid residues, of these 4 were selected based on their position on the pCP312R sequence, on the amino end, the center of the sequence and on the carboxyl end of the sequence. A graph of antigenic propensity against sequence position is shown in figure 2;

**Fig 2:**
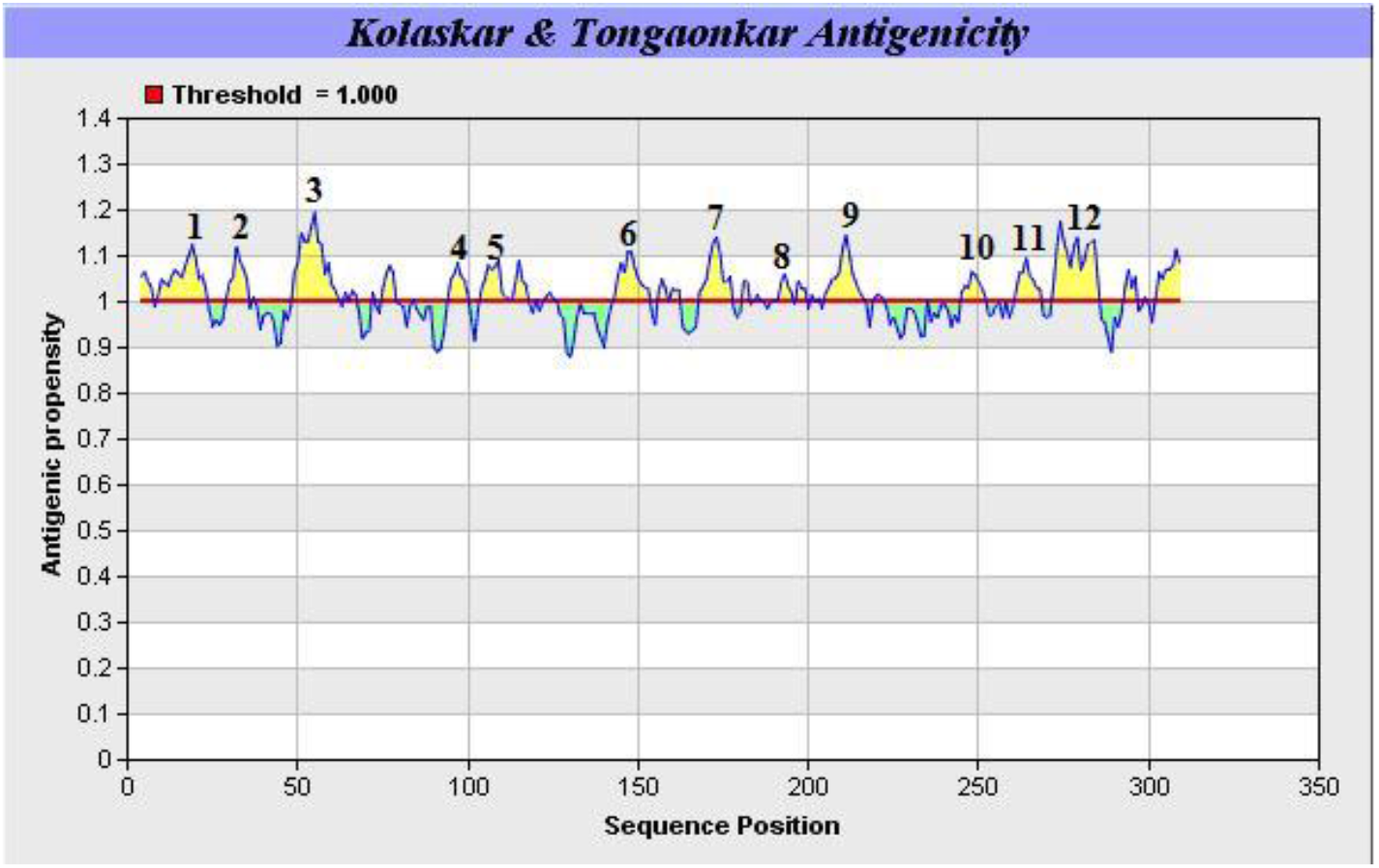
Antigenic propensity of pCP312R sequence as predicted by Kolaskar and Tongaonkar method. The X-axis represents sequence positions which are predicted to be highly antigenic while the Y-axis represents antigenic propensity. The average antigenic propensity of the sequence is 1.017, while the minimum and maximum antigenic propensities are 0.878 and 1.196 respectively.

The sequences of the predicted antigenic peaks, their lengths and position on the pCP312R sequence are shown in table 3;

**Table 3:**
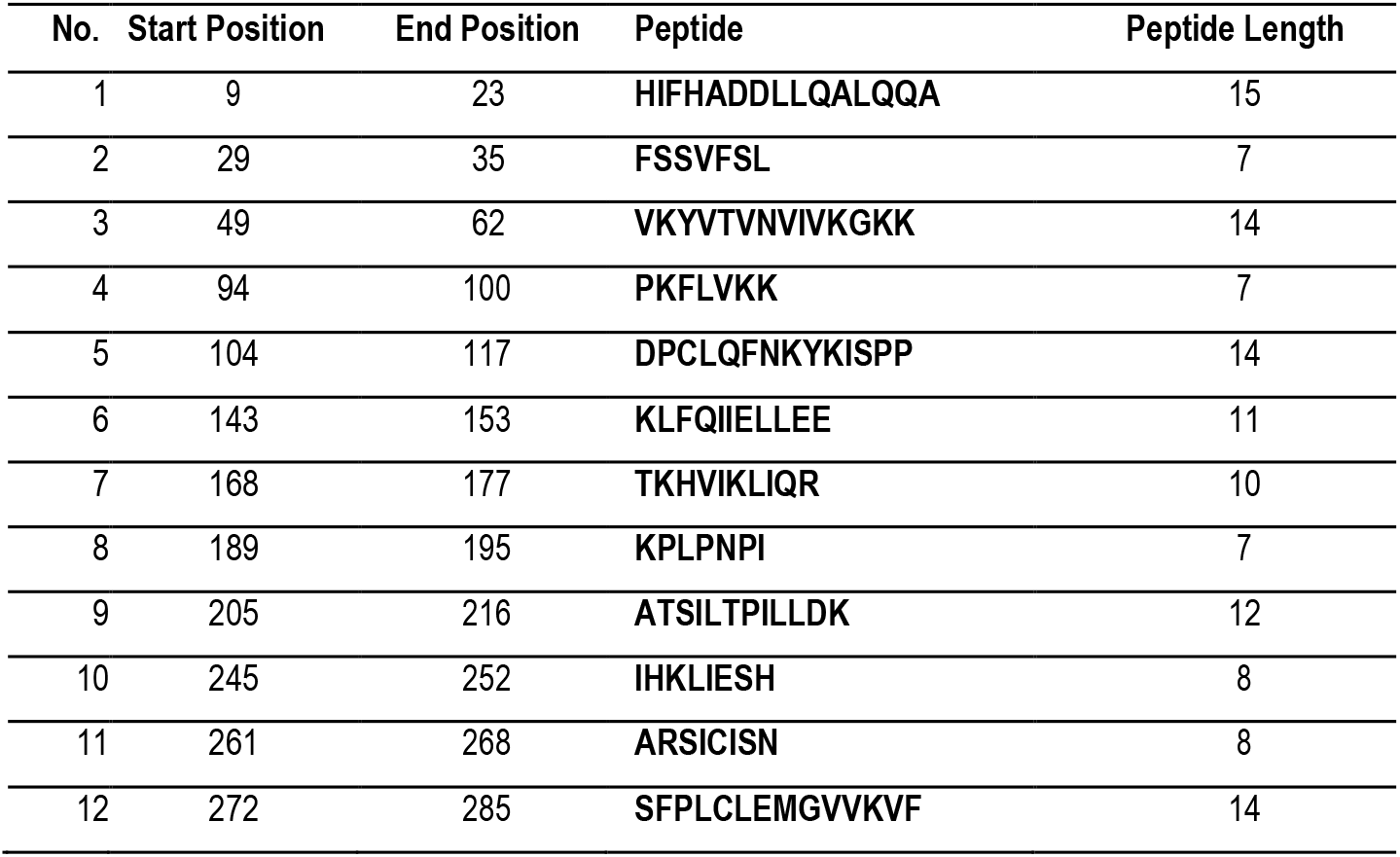
Properties of the antigenic regions of pCP312R sequence

Peptides number 1, 3, 5 and 12 were selected, synthesised and used in immunisation of rabbits

### Immune response of rabbits to synthetic peptides

Immune response of rabbits to each of the peptides was compared against each other and against pre - immune serum, used as a negative control. Peptide cpr1 gave the highest response followed by cpr2 then cpr3 and cpr4 gave the same response when optical density readings were compared from serum collected at day 52. The peptide from p54 used as a positive control in this study gave optical density values higher than all the peptides studied (OD 3.5), as shown in figure 3 below;

**Fig 3:**
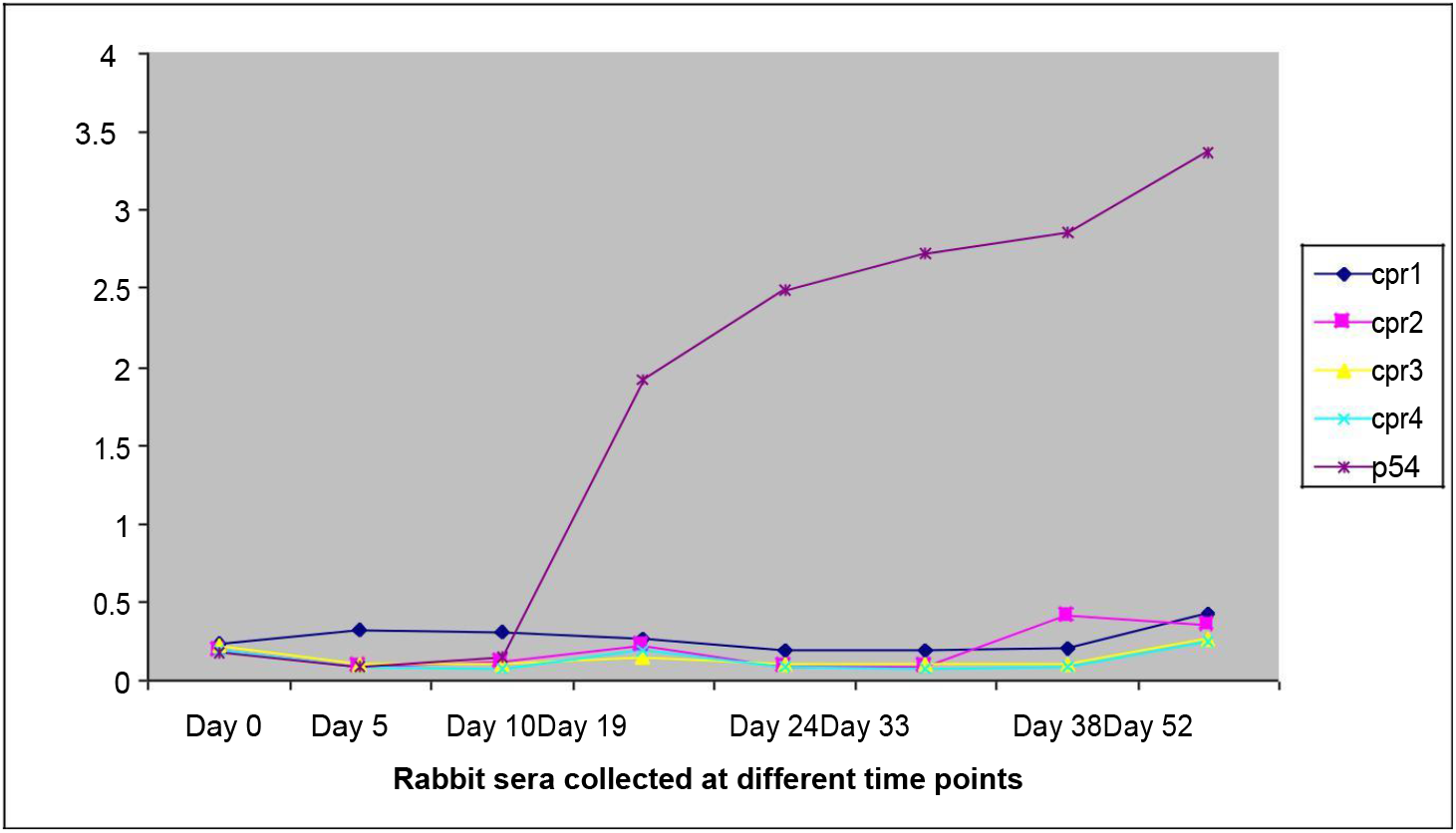
Graph showing rabbit immune response against synthetic peptides. Rabbit serum was collected from each group of rabbits and analysed by ELISA at a dilution of 1:20,000 (primary antibody) and 1:10,000 (secondary antibody). The graph shows analysis of pre immune serum as well as immune serum collected at seven different time points from day 0 to day 52.

### Results of the Immunohistochemistry evaluation using polyclonal anti-peptide antibodies

Immunohistochemistry using antibodies against the peptides was then performed on ASFV positive tissue, staining was observed in macrophages as well as within interstitial tissue. Antibodies against peptides cpr1, cpr2, cpr3 and cpr4 demonstrated specific staining of macrophages, and staining in interstitial when used at a dilution of 1:200. Pre-immune serum was used as a negative control and did not stain ASFV positive tissue at 1:200 dilution, also tissue from an uninfected pig was used as a negative control and stained with anti - peptide antibodies and did not stain, while anti p54 antibodies was used as a positive control at a dilution of 1:200 was able to stain macrophages specifically. Representative results obtained are shown in Figures 4 and 5. Figure 4 shows Photomicrographs of positive immunostaining of ASFV while Figure 5 shows comparison of positive immunostaining of ASFV amongst the four different anti-peptide polyclonal antibodies;

**Fig 4:**
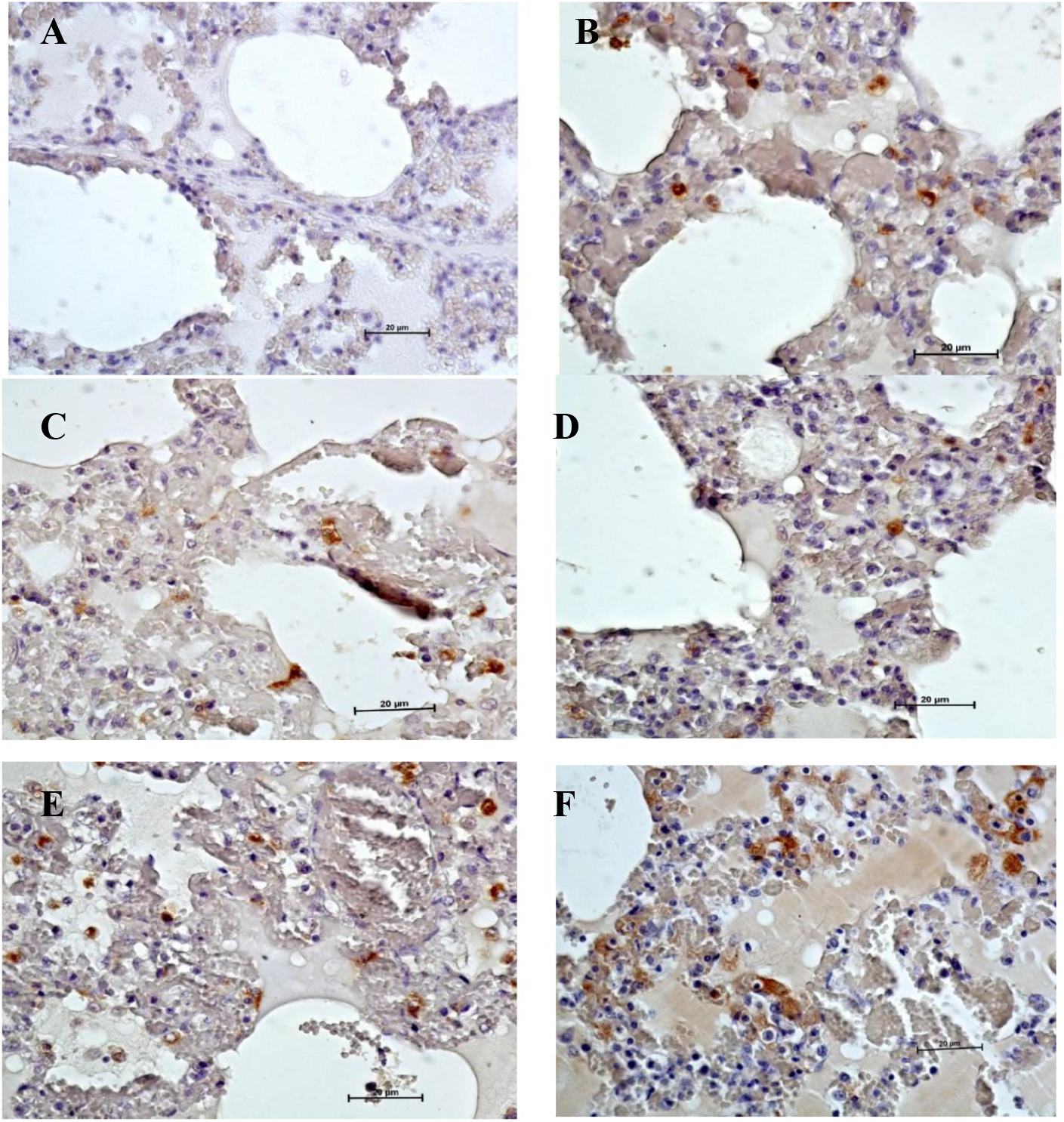
Photomicrographs showing positive immunostaining of ASFV in different sections of the same lung tissue of an affected pig using antibodies against synthetic peptides. A is a negative control with pre-immune serum; B is a staining with polyclonal anti-cpr1, C is staining with anti-cpr2, D is staining with anti-cpr3, E is staining with anti-cpr4 and F is staining with anti-p54. ASFV infected macrophages and lysed cells are stained brown, all anti-peptide sera demonstrated specific staining. Antibodies were diluted at 1:200. Immunostaining was done with Histofine SAB-PO immunohistochemistry kit (Nichirei, Tokyo, Japan), with DAB reagent (Dako, DAB kit Japan) and counter stained with Mayer’s Haematoxylin X400.

**Fig 5:**
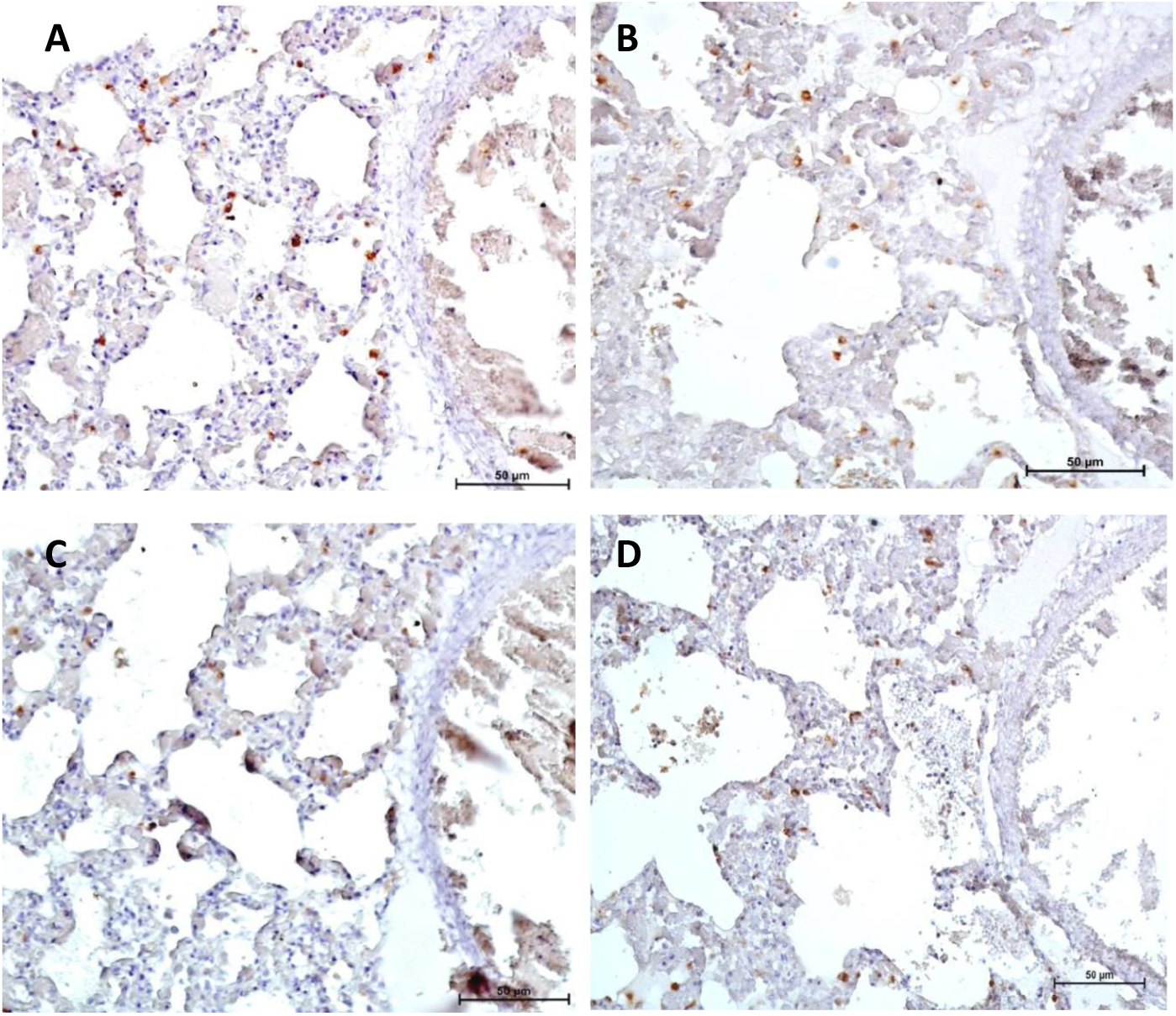
Photomicrographs showing comparison of positive immunostaining of ASFV amongst the four different anti-peptide polyclonal antibodies in different sections of the same lung tissue of an infected pig. A is a staining with polyclonal anti-cpr1, B is staining with anti-cpr2, C is staining with anti-cpr3, D is staining with anti-cpr4. ASFV infected macrophages and lysed cells are stained brown, anti-cpr1, anti-cpr2 and anti-cpr4 gave a staining score of +++ while anti-cpr3 gave a score of ++. Antibodies were diluted at 1:200. Immunostaining was done with Histofine SAB-PO immunohistochemistry kit (Nichirei, Tokyo, Japan), with DAB reagent (Dako, DAB kit Japan) and counter stained with Mayer’s Haematoxylin X200.

Comparison of the staining ability of the four anti-peptide antibodies was performed and number of stained macrophages in a similar area of different sections of the same tissue was used to obtain a score, 1 −10 stained cells was given a score of +, 11-20 a score of ++, and 21-30 a score of +++. Anti-cpr1, cpr2 and cpr4 gave a score of +++ while anti-cpr3 antibodies gave a score of ++. Figure 5 shows the representative results of the comparison;

### Results of evaluation of diagnostic potential of pCP312R peptides by ELISA

Out of the 540 porcine field sera samples collected and screened using OIE recommended ELISA and PCR tests, 20 samples were found to be positive by both ELISA and PCR. These together with 25 negative samples were selected and used for evaluation of the diagnostic potential of pCP312R peptides. The diagnostic sensitivity and specificity of each ELISA was calculated at 95% CI and 5% allowable error. The differences in optical densities between positive and negative samples were small therefore the results were categorised into positive and negative, with no intermediates.

Synthetic peptide CP1 gave a diagnostic sensitivity of 55% (95% CI, 0.3421-0.7418) and specificity of 96% (95% CI, 0.8046-0.9929), peptide CP2 gave a diagnostic sensitivity of 100% (95% CI, 0.8389-1) and specificity of 52% (95% CI, 0.335-0.6997), Synthetic peptide CP3 gave diagnostic sensitivity of 95% (95% CI, 76.39-99.11) and specificity of 88% (95% CI, 70.04-95.83), peptide CP4 gave a diagnostic sensitivity of 90% (95% CI: 0.699-0.9721) and specificity of 76% (95% CI: 0.5657-0.885) and positive control peptide p54 gave a diagnostic sensitivity of 100% (95% CI, 0.8389-1) and specificity of 56% (95% CI, 0.3707-0.7333). The diagnostic sensitivity and specificity of the OIE ELISA is 98.8% and 87.8% respectively. The results are summarised in the table 4;

**Table 4:** Summary of the results of evaluation of the different synthetic peptides by ELISA

| Peptides evaluated by ELISA |  |  |  |  |  |
| --- | --- | --- | --- | --- | --- |
|  | CP1 | CP2 | CP3 | CP4 | P54 |
| **Sensitivity** | **0.55**[0.342- 0.742] | **1.000**[0.804-0.997] | **0.95**[0.764-0.991] | **0.9**[0.699-0.972] | **1.000**[0.804- 0.997] |
| **Specificity** | **0.96**[0.805- 0.993] | **0.52**[0.335-0.7] | **0.88**[0.7-0.958] | **0.76**[0.566- 0.885] | **0.56**[0.371-0.733] |
| **PPV** | **0.917**[0.646-0.985] | **0.625**[0.453- 0.771] | **0.864**[0.667- 0.953] | **0.75**[0.551-0.88] | **0.645**[0.469- 0.789] |
| **NPV** | **0.727** [0.558-0.849] | **1.000**[0.725- 1.000] | **0.957**[0.79-0.992] | **0.905**[0.711- 0.973] | **1.000**[0.74-1.000] |
| **LR+** | **13.75**[1.935- 97.699] | **2.033**[1.344-3.074] | **7.917**[2.726- 22.994] | **3.75**[1.839-7.648] | **2.217**[1.417- 3.469] |
| **LR-** | **0.469**[0.287- 0.766] | **0.047**[0.003- 0.744] | **0.057**[0.008-0.386] | **0.132**[0.035- 0.499] | **0.044**[0.003- 0.688] |

Key: PPV is **Positive predictive value**, NPV is **Negative Positive predictive value**, LR+ is **Positive likelihood ratio**, and LR-is **Negative likelihood ratio**.

## Discussion

The aim of this study was to contribute knowledge towards the identification and evaluation of new candidate molecules for the sero-diagnosis of African swine fever. African swine fever is endemic in Uganda with outbreaks reported throughout the year (Atuhaire *et al*., 2013). To date there is no treatment or vaccine available for African swine fever, therefore control of the disease relies on accurate diagnostic procedures, as the first step followed by quarantine of infected animals. Diagnosis of ASF in Uganda mainly relies on the clinical signs and post-mortem lesions. However, the clinical signs and lesions are not pathognomonic for ASF. This study successfully investigated the immunogenicity of synthetic peptides mimicking an ASFV putative protein pCP312R and evaluated the diagnostic potential of an ELISA test based on these peptides; in addition this study evaluated the use of antibodies produced against these peptides, in diagnosis of ASF by immunohistochemistry.

In silico prediction of antigenic epitopes using the method described by Kolaskar and Tongaonkar predicted an average propensity of 1.017 which means that all amino acid residues having a propensity value above this average value were considered potentially antigenic. The selected peptides all had propensity values above average thus suggesting that they were antigenic. When antigenic propensity values of ASFV pCP312R sequence were compared with values of other ASFV proteins, the antigenic propensity values were similar to those of known ASFV diagnostic proteins vp73 and p54 (see supplementary material). The average antigenic propensity of pCP312R was 1.017 while that for vp73 and p54 were 1.026 and 1.031 respectively. Comparison of the highest peaks, that is maximum antigenic propensity also showed that pCP312R has almost the same maximum antigenic propensity (1.196) as vp73 (1.197) though slightly lower than p54 (1.233) (see supplementary material). So by in silico comparison alone pCP312R was shown to have properties that suggest it has highly antigenic regions. It should however be noted that this method of antigenic prediction is known to be 75% accurate and therefore has a 25% chance of not predicting the right antigenic sites although it is more accurate than most methods.

The immune response of rabbits to these selected synthetic peptides was then studied using an indirect ELISA. The antibody titers varied for each of the peptides although all the four peptides derived from the pCP312R sequence gave relatively low optical density values as compared to the peptide derived from the p54 sequence. This low optical density readings showed that even if these peptides do elicit an immune response, the antibody titers are lower than those obtained from immunisation with p54 peptide which was being used as a control for the immunisations. This p54 peptide elicited a higher immune response than the pCP312R derived peptides. This is in agreement with previous studies that have shown that P54 is highly immunogenic (Barderas *et al*., 2001; Rodriguez *et al*., 1994). Amongst the pCP312R peptides, cpr1 and cpr2 gave higher optical density readings than the cpr3 and cpr4. Peptides cpr1 and cpr2 were selected from the amino end of the protein sequence thus suggesting that immunogenicity of pCP312R may be dependent on location of antigen on the protein sequence.

This study further evaluated the diagnostic potential by immunohistochemistry, of polyclonal antibodies raised against these synthetic peptides. Analysis of the antibodies raised against each of the peptides revealed ability to detect viral antigen in ASF positive tissue although with varying levels of staining intensity with DAB (chromogen). This confirms that the peptides are immunogenic and optimal staining was obtained at a dilution of 1:200, which means that the antibodies are present but in low concentration. However this dilution is prone to non specific staining (background). Improving antibody titers may however have been done by increasing immunisation dosage, optimisation of most suitable peptide conjugate protein and combination of adjuvants, and using a different animal model for the immunisation study. This in addition to using purified polyclonal IgG and monoclonal antibodies would probably increase the concentration and titers of the anti -peptide antibodies. Despite not having high antibody titers, the anti-peptide sera contained specific antibodies.

Analysis of sera raised against pCP312R derived peptides gave anti-cpr1, anti-cpr2 and anti-cpr4 antibodies with better staining specificity of virus antigens in infected tissue than anti-cpr3. Anti-p54 antibodies, used as a control also demonstrated high staining ability, amongst all the peptides studied. Previous studies have performed immunohistochemistry for ASF using monoclonal anti-vp73 antibody (Fernández *et al*., 2007; Pe’rez *et al*., 1994).

This study has shown for the first time that polyclonal antibodies collected from rabbits, immunised with synthetic peptides mimicking antigenic epitopes of ASFV protein are a promising candidate for immunohistochemical diagnosis of ASF. This study presents a fairly affordable method that is comparable to currently available immunohistochemistry tests and does not require laborious and expensive virus isolation to obtain proteins to be used in production of diagnostic sera. We have further shown that the ASFV pCP312R protein is immunogenic and should be extensively studied as a whole protein, probably in recombinant form. It also shows that antibodies raised against linear epitopes can be used in immunohistochemical detection of ASFV. Peptides have previously been used in ASF vaccine development studies (Ivanov *et al*., 2011) but to date, no published study has looked at the use of peptides in diagnosis of ASF.

This study confirms that this method of prediction of antigenic epitopes is accurate, since the peptides selected using this method were able to elicit an immune response in rabbits and the antibodies obtained were also able to detect ASFV antigen in infected pig tissues. However not all predicted peptides were used in this study, thus potentially missing out on some highly immunogenic peptides.

ASFV peptides were additionally evaluated for use in diagnostic ELISA, using serum samples of known infection status. The peptides evaluated in this study demonstrated varying levels of accuracy when used in indirect ELISA tests by recording diagnostic sensitivities as low as 55%, as shown by peptide CP1 (sensitivity of 55% (95% CI, 0.3421-0.7418) and specificity of 96% (95% CI, 0.8046-0.9929)) and diagnostic specificities as low as 52% for peptide CP2 (sensitivity of 100% (95% CI, 0.8389-1) and specificity of 52% (95% CI, 0.335-0.6997)). CP3 gave a diagnostic sensitivity of 95% (95% CI, 76.39-99.11) and specificity of 88% (95% CI, 70.04-95.83), CP4 gave a diagnostic sensitivity of 90% (95% CI: 0.699-0.9721) and specificity of 76% (95% CI: 0.5657-0.885) and p54 gave a diagnostic sensitivity of 100% (95% CI, 0.8389-1) and specificity of 56% (95% CI, 0.3707-0.7333).

Notably however peptides CP2 and p54 (control) had diagnostic sensitivity’s of 100% thus suggesting that the two peptides may be further evaluated for use in other serological tests for mass screening of pig herds, in order to detect exposure to virus antigens before confirming infection with a more accurate method. The peptide cp3 gave a specificity of 88% which is slightly higher than the specificity of the OIE ELISA (87.8%). This combined with a sensitivity of 95% makes this peptide a promising candidate for serodiagnosis of ASF. The performance of peptide ELISA’s was however generally weaker than previously described ASF diagnostic ELISA using semi purified virus antigens (OIE 2008 and those using ASFV recombinant proteins (Barderas *et al*., 2000, Gallardo *et al*., 2006, Gallardo *et al*., 2011). It must be noted that with immunohistochemistry the results are promising but it is not the case with ELISA, this may probably be due to a number of reasons; very little peptide binding on the ELISA plate wells, since coating was done with unconjugated peptides, unsuitable coating buffer used and quality of plates used. It must also be noted that the four synthetic peptides evaluated here are part of the same protein sequence and despite them not being very promising individually, their performance when combined may be better since these peptides have demonstrated different strengths. Limitation of presentation of results to either positive or negative, with no intermediates, may have also contributed to low sensitivity and specificity values for some of the peptides.

The peptides evaluated in this study have shown promise as candidate antigens for diagnosis of ASF. They could be explored for use in simple tests that can be moved closer to end users for example rapid immunohistochemical tests, card agglutination tests, and immunoblotting, for rapid detection of ASFV.

## Conclusion

This study presents the first time synthetic peptides have been evaluated for serodiagnosis of African swine fever in domestic pigs. The study successfully predicted and designed peptides with varying levels of immunogenicity. This study in addition showed that there is potential for use of polyclonal anti-peptide serum in the diagnosis of ASF using immunohistochemistry. This study on the other hand did not give conclusive results for the use of synthetic peptides as candidates for serodiagnosis of ASF using ELISA.

### Authors’ contributions

SO contributed to the conception of the idea, design, data collection, laboratory studies, drafting and writing of the manuscript. MA contributed to laboratory studies and manuscript preparation. DKA contributed to laboratory studies, data analysis and drafting of the manuscript. MK contributed to data collection, data analysis and writing of the manuscript. PKM contributed to data collection, laboratory studies and writing of the manuscript. JBO contributed to conception of the idea, data collection and writing of the manuscript. WO contributed to conception of the idea, design and writing of the manuscript. LO contributed to conception of the idea, design, data collection and writing of the manuscript. All read and approved the manuscript.

### Authors’ information

SO is a MSc holder with interest in African swine fever and Foot and mouth disease virus diagnostics. MA is a PhD holder working on African swine fever in Uganda, PKM is a PhD student working on human African Trypanosomiasis, MK is a MSc student working on Paratuberculosis. DKA is a PhD holder with expertise in Veterinary diagnostics, JBO is a PhD holder with expertise in Veterinary diagnostics. WO is a PhD holder and has researched widely in the livestock sector. LO is a PhD holder and a Professor of Veterinary Pathology with vast experience on diagnosis of livestock diseases.

## Acknowledgement

This study was funded by the Millennium Science Initiative through a grant to Professor Ojok Lonzy, Dr. William Olaho-Mukani, and Dr. JB.Okuni of the Appropriate Animal Diagnostic Technologies project under the Uganda National Council of Science and Technology.

