## Supplemental figures for "Serodiagnostic potential of synthetic peptides derived from African swine fever virus putative protein pCP312R" (from another version of this preprint: version 1, 2019-01-07)

**Kolaskar & Tongaonkar Antigenicity for p54 protein**

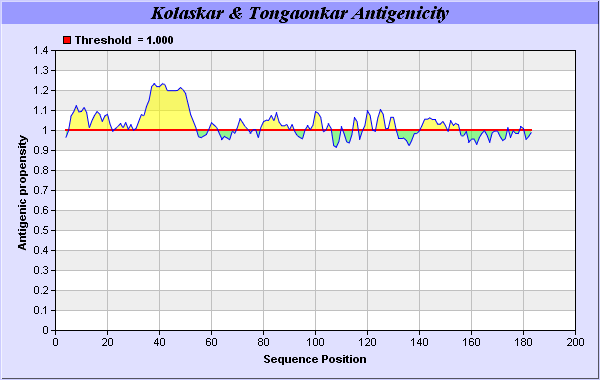

**Average:** 1.031   **Minimum:** 0.917   **Maximum:** 1.233   **Threshold:** 1.000

Top of Form

**Sequence:**

        1 MDSEFFQPVY PRHYGECLSS TPTPSFFSTH MYTILIAIVV LIIIIIVLIY LFSSRKKKAA 
       61 AAIEEEDIQF INPYQDQQWA GVPPQPGIAK PAGATTGSVS KPVMDRPITN NPVMDRPVTN 
      121 HPVTDRLVTD KLGMATGEPA AASAPAHSAE PYTTVTTQNT ASQTMSAIEN LRQRSTYTHK 
      181 DLENSL

### Kolaskar & Tongaonkar Antigenicity for vp73 protein

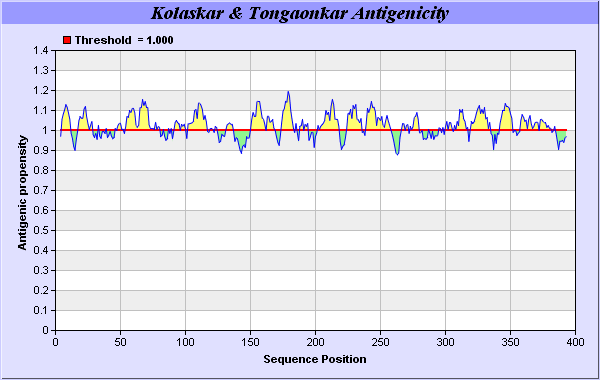

**Average:** 1.026   **Minimum:** 0.879   **Maximum:** 1.197   **Threshold:** 1.000

**Sequence:**

        1 MASGGAFCLI ANDGKADKII LAQDLLNSRI SNIKNVNKSY GKPDPEPTLS QIEETHLVHF 
       61 NAHFKPYVPV GFEYNKVRPH TGTPTLGNKL TFGIPQYGDF FHDMVGHHIL GACHSSWQDA 
      121 PIQGTAQMGA HGQLQTFPRN GYDWDNQTPL EGAVYTLVDP FGRPIVPGTK NAYRNLVYYC 
    181 EYPGERLYEN VRFDVNGNSL DEYSSDVTTL VRKFCIPGDK MTGYKHLVGQ EVSVEGTSGP 
  241 LLCNIHDLHK PHQSKPILTD ENDTQRTCSH TNPKFLSQHF PENSHNIQTA GKQDITPITD 
  301 ATYLDIRRNV HYSCNGPQTP KYYQPPLALW IKLRFWFNEN VNLAIPSVSI PFGERFITIK 
      361  LASQKDLVNE FPGLFIRQSR FIPGRPSRRN IRFKPW
